# Constrained hypermutation and absence of TERT promoter mutations in Lynch syndrome-associated urothelial cancer

**DOI:** 10.1101/2024.11.27.624238

**Authors:** Jussi Nikkola, Lauri Ryyppö, Juuso Vuorinen, Lauri Moilanen, Maarit Ahtiainen, Kirsi Pylvänäinen, Hanna Selin, Tuomo Virtanen, Matti Nykter, Thea Veitonmäki, Jukka-Pekka Mecklin, Toni T. Seppälä, Matti Annala

## Abstract

Lynch syndrome (LS) is a hereditary condition characterized by defective DNA mismatch repair (MMR) and high incidence of several cancers, including urothelial cancers (UC) of the upper urinary tract and bladder. We set out to study the somatic landscape of LS-associated urothelial cancer (LS-UC) by analyzing 41 surgical tumor samples and 3 urine DNA samples from 34 LS-UC patients. We show that telomerase reverse transcriptase (*TERT*) promoter mutations found in 83% of sporadic UC are almost completely absent (5%) in LS-UC (p < 0.00001). Instead, all LS-UC carried a 5-methylcytosine deamination (CG>TG) and microsatellite instability driven mutation landscape characterized by highly frequent *ARID1A* (82%), *FGFR3* (80%), and KMT2D (78%) mutations, as well as preferential usage of CG>TG mutation hotspots. We propose that the scarcity of *TERT* promoter mutations in LS-UC is due to inability to create the necessary GABP binding motif (5’-GGAA) through CG>TG mutation or microsatellite instability. Our data shows that LS-UC represents a disease entity with unique genomic characteristics.

## Main Text

Lynch syndrome (LS) is a dominantly inherited cancer predisposition syndrome characterized by inactivating germline mutations in DNA mismatch repair (MMR) genes *MSH2, MSH6, MLH1, PMS2*, or the *EPCAM* gene which regulates *MSH2* expression [1]. Affected patients suffer from a high incidence of colon and endometrial cancer, but also a 10 - 25% lifetime risk of urothelial cancer, particularly in the upper urinary tract [2]. LS-associated cancers typically obtain an inactivating second hit mutation or epigenetic change to the predisposing gene as an early event in tumorigenesis [3]. The second hit disrupts DNA repair resulting in accelerated mutagenesis, with particular enrichment of CG>TG mutations arising from spontaneous 5-methylcytosine deamination, and somatic changes to microsatellite lengths [4].

The most common somatic genomic alteration in sporadic urothelial cancers (UC) is mutation of the telomerase reverse transcriptase (*TERT)* promoter, occurring at one of two mutation hotspots (−124C>T or -146C>T) in 70 - 80% of patients [5]. These mutations increase *TERT* expression by creating a new GA-binding protein (GABP) transcription factor binding site with the core motif 5’-GGAA-3’, and are an early event in urothelial tumorigenesis [6]. TERT plays a key role in telomere elongation, and its overexpression in urothelial cancer enhances this activity [7]. Sporadic UC tumors also harbor recurrent mutations in *FGFR3, TP53, PIK3CA, KDM6A, ERBB2, ERBB3*, and *STAG2* [8]. A study by Donahue et al. provided early evidence that Lynch syndrome-associated urothelial cancers (LS-UC) carry more mutations in several genes including *FGFR3* and *ARID1A*, but did not investigate the *TERT* promoter region and analyzed only a small cohort consisting solely of upper tract urothelial tumors (UTUC) [9].

To investigate the genomic drivers of bladder and upper tract urothelial cancer in LS patients, we searched all UC diagnoses for patients in the Finnish Lynch Syndrome Registry, and collected 41 UC surgical tissue samples accrued to Finnish biobanks between April 1987 - June 2022 from a total of 32 LS patients (11 bladder cancer, 21 UTUC). We also analyzed 125 tumor DNA positive urine samples collected between January 2021 and January 2024 from 122 sporadic UC and 3 upper tract LS-UC patients recruited on an all-comers basis to the UROLIB study at the Tampere University Hospital (**Supplementary Table 1**). All 34 LS-UC patients (one patient provided tissue and urine) were confirmed carriers of a pathogenic germline MMR variant (16 *MSH2*, 13 *MLH1*, 5 *MSH6*). LS-UC cases were more often UTUC (68% vs 14%, p < 0.0001; of which ureteral 67% vs 29%, p = 0.030), young (median 62 vs 74 years, p < 0.0001), and female (48% vs 23%, p = 0.0046) compared to sporadic UC (**Table 1**), consistent with prior literature [10].

**Table 1.** Clinical characteristics of the Lynch syndrome-associated urothelial cancer (LS-UC) and sporadic urothelial cancer (UC) patients at the time of tumor sample collection (surgical tissue or urine tumor DNA for LS-UC, urine tumor DNA for the sporadic UC cohort).

| Characteristic | LS-UC tumors (n = 44) | Sporadic UC tumors (n = 122) |
| --- | --- | --- |
| **Tumor subtype** |  |  |
| Non-muscle invasive bladder cancer | 11 (25%) | 83 (68%) |
| pTa low grade | 1 (2%) | 46 (38%) |
| pTa high grade | 5 (11%) | 23 (19%) |
| pT1 | 5 (11%) | 13 (11%) |
| pTis | 0 (0%) | 1 (1%) |
| Muscle invasive bladder cancer | 3 (7%) | 22 (18%) |
| pT2 | 2 (5%) | 17 (14%) |
| pT3 | 1 (2%) | 1 (1%) |
| pT4 | 0 (0%) | 4 (3%) |
| Upper tract urothelial carcinoma | 30 (68%) | 17 (14%) |
| pTa low grade | 3 (7%) | 1 (1%) |
| pTa high grade | 7 (16%) | 2 (2%) |
| pTis | 1 (2%) | 0 (0%) |
| pT1 | 6 (14%) | 1 (1%) |
| pT2 | 8 (18%) | 2 (2%) |
| pT3 | 4 (9%) | 8 (7%) |
| pT4 | 1 (2%) | 3 (2%) |
| **Sex** |  |  |
| Male | 23 (52%) | 94 (77%) |
| Female | 21 (48%) | 28 (23%) |
| **Age** |  |  |
| 39 - 49 years | 5 (11%) | 3 (2%) |
| 50 - 59 years | 8 (18%) | 7 (6%) |
| 60 - 69 years | 18 (41%) | 25 (20%) |
| 70 - 79 years | 10 (23%) | 53 (43%) |
| 80 - 95 years | 3 (7%) | 34 (28%) |
| **Hematuria** |  |  |
| Macroscopic hematuria | 24 (55%) | 104 (85%) |
| Microscopic hematuria | 0 (0%) | 1 (1%) |
| No hematuria | 13 (30%) | 17 (14%) |
| Unknown | 7 (16%) | 0 (0%) |

All tissue and urine samples were analyzed using the UroScout assay, a hybridization capture panel targeting 25 urothelial cancer associated genes including the MMR genes *MSH2, MSH6, MLH1*, and *PMS2* [11] (**Figure 1a, Supplementary Table 2**). The pathogenic germline MMR variant was detected by UroScout in all LS-UC patients, and a somatic second hit to the same gene was detected in 28/34 cases (**Figure 1a, Supplementary Table 1**). Immunohistochemistry confirmed loss of the relevant MMR protein in all LS-UC tumor samples (**Supplementary Figure 1**).

**Figure 1.**
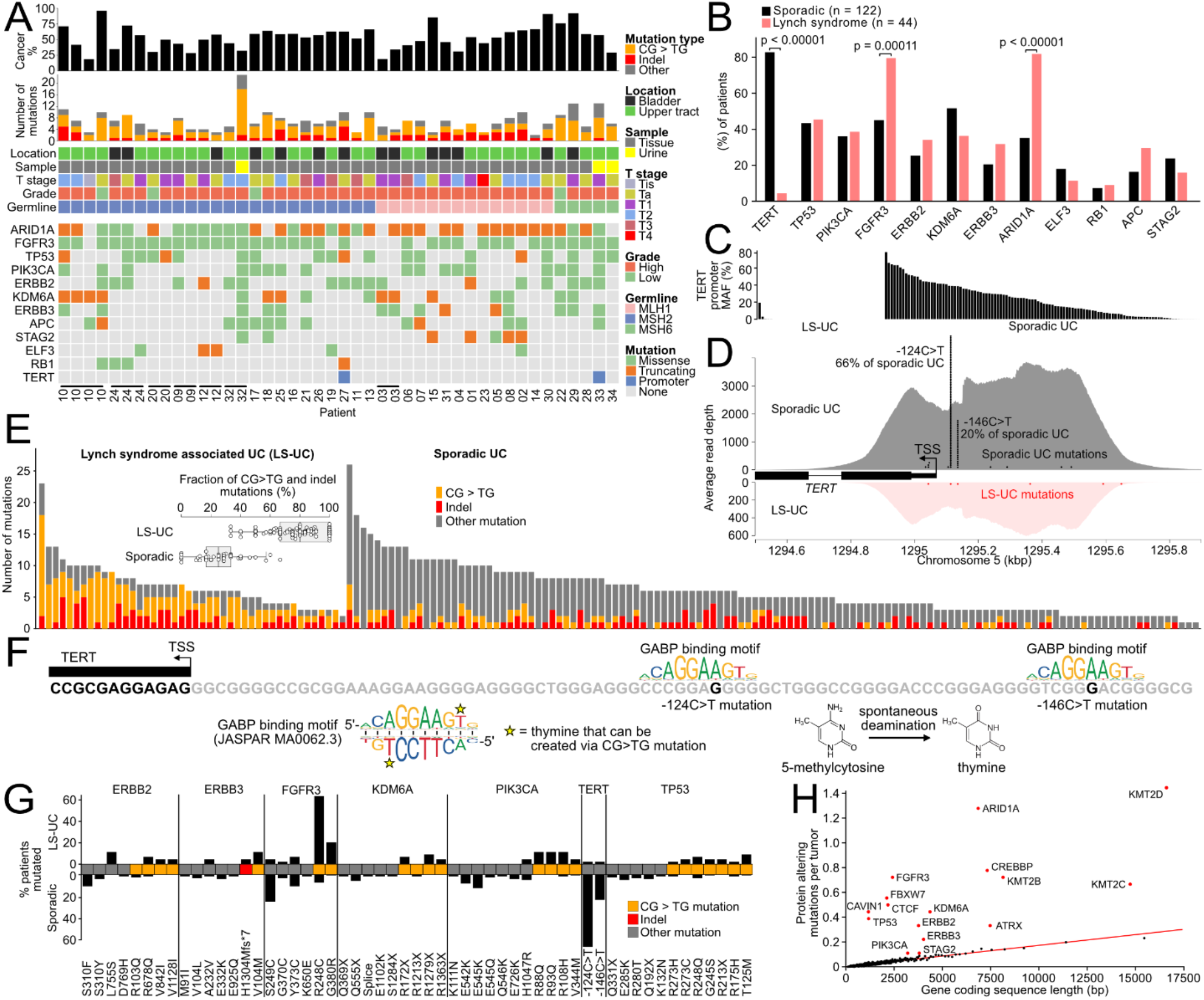
Somatic genomic landscape of Lynch syndrome-associated urothelial cancer (LS-UC). (**a**) Cohort overview and somatic mutation landscape. Same-patient tissue samples are indicated with horizontal lines underneath. Patients are sorted by germline defect. (**b**) Comparison of driver gene mutation frequency between sporadic and LS-UC tumors. (**c**) Highest measured allele fraction for the two primary TERT promoter mutations in each sporadic and LS-UC sample. (**d**) Visualization of sequencing depth and detected mutations across the TERT promoter region in sporadic and LS-UC. (**e**) Enrichment of CG>TG and indel type somatic mutations in LS-UC relative to sporadic UC. (**f**) Visualization of the *TERT* upstream region and the GABP binding motifs created by the two most common TERT promoter region mutations -124C>T and -146C>T, neither of which is formed through CG>TG 5-methylcytosine deamination. In general, CG>TG mutations cannot produce the core 5’-GGAA GABP binding motif as it contains no TG dinucleotides. (**g**) Mutation frequency at recurrent mutation hotspots in sporadic and LS-UC. CG>TG, indel, and other mutations are indicated using color. (**h**) Relationship between gene length and the frequency of protein altering mutations in LS-UC tumors based on whole exome sequencing. Genes were sorted by coding sequence (CDS) length, and then combined into groups of 50 genes (dots). Mutation frequencies of known UC driver genes are indicated as large red dots.

Telomerase reverse transcriptase (*TERT*) gene promoter mutations were detected in 101/122 (83%) sporadic UC tumors, but only 2/44 (5%) LS-UC tumors (p < 0.0001, Fisher’s exact test) (**Figure 1b-c**). We found no alternative mutations or rearrangements in LS-UC tumors producing a GABP binding motif, nor other recurrent mutations or rearrangements within the 500 bp *TERT* upstream region covered by our panel (**Figure 1d**). Conversely, *FGFR3* and *ARID1A* were mutated in ≥80% of LS-UC cases, an elevated rate relative to sporadic UC (**Figure 1b)**. We also compared the LS-UC mutation frequencies against a published cohort of 2463 sporadic UC tissues, and verified that the differences in driver gene mutation frequency were not simply explained by the anatomic location, sex, or age of LS-UC cases (**Supplementary Figures 2 - 5**). As further confirmation for the scarcity of *TERT* promoter mutations in LS-UC, we noted that a previous independent UTUC study by Fujii et al. found no *TERT* promoter mutations in the seven LS-UTUC tumors in their cohort [12]. More generally, an enrichment of CG>TG and insertion/deletion type mutations was found in all LS-UC cases relative to sporadic UC (75% vs 22% of all mutations, p < 0.0001), implicating MMR defects as the primary driver of LS-UC tumorigenesis (**Figure 1e, Supplementary Table 3**). The mutation landscapes of LS-associated bladder and upper tract tumors showed no differences (**Supplementary Figure 6**).

We propose that *TERT* promoter mutations are absent in LS-UC because the core GABP binding motif 5’-GGAA-3’ cannot be created via CG>TG mutation or microsatellite instability, resulting in tumorigenesis via an alternative set of somatic driver alterations (**Figure 1f, Supplementary Figure 7**). Constrained evolution of LS-UC was evident in the mutation landscapes of other driver genes such as *FGFR3, PIK3CA*, and *TP53*, strongly favoring well-characterized CG>TG mutation hotspots (84% vs 9% of mutations at recurrently mutated sites, p < 0.0001) instead of mutation hotspots characteristic to sporadic UC (**Figure 1g**). Whole exome sequencing of 18 *TERT* promoter mutation negative LS-UC tumors found 5 (28%) cases with a mutation in *ATRX*, a gene whose inactivation is linked to telomere lengthening [13]. Exome sequencing also revealed recurrent LS-UC mutations in genes *KMT2D, FBXW7, CREBBP, CTCF*, and *KMT2C*, each having a higher mutation frequency than expected based on CDS length (**Figure 1h**). *ARID1A* inactivation has been shown to increase *TERT* expression in human cells [14].

In conclusion, we show that LS-UC is characterized by a unique mutation landscape, including a striking absence of *TERT* promoter mutations. Our proposed explanation that *TERT* mutations are absent in LS-UC due to constrained hypermutation is supported by similar findings in other UC driver genes, but we cannot rule out alternative explanations such as rapid tumorigenesis resulting in lower mitotic age and reduced need for *TERT* overexpression. The absence of *TERT* promoter mutations in LS-UC is relevant for diagnostic tests relying on these recurrent mutations. While immunotherapy remains the primary treatment option for dMMR cancer patients with excellent outcomes [15], the higher prevalence of FGFR mutations in LS-UC suggests that FGFR inhibitors could be investigated as a therapeutic option for this patient population even for localized stages.

### Financial disclosures

Matti Annala and Toni Seppälä certify that all conflicts of interest, including specific financial interests and relationships and affiliations relevant to the subject matter or materials discussed in the manuscript (e.g. employment/affiliation, grants or funding, consultancies, honoraria, stock ownership or options, expert testimony, royalties, or patents filed, received, or pending), are the following: Juuso Vuorinen is a co-founder of Fluivia. Toni Seppälä declares consultation fees from Tillots Pharma, Nouscom and Mehiläinen, is CEO and co-owner of Healthfund Finland, and a clinical advisory board member and minor shareholder of Lynsight. Matti Annala is CEO and co-founder of Fluivia. The remaining authors have nothing to disclose.

## Acknowledgements

The authors acknowledge the Biocenter Finland (BF) and Tampere Genomics Facility for their service. The authors wish to acknowledge CSC – IT Center for Science, Finland, for computational resources. The study benefited from samples from the Helsinki, Tampere, Auria, Central Finland, Eastern Finland, and Northern Finland biobanks. We are grateful to all participating patients and their families.

## References

[1] Ligtenberg MJL, Kuiper RP, Chan TL, Goossens M, Hebeda KM, Voorendt M, et al. Heritable somatic methylation and inactivation of MSH2 in families with Lynch syndrome due to deletion of the 3′ exons of TACSTD1. Nat Genet 2008;41:112–7.

[2] Møller P, Seppälä TT, Bernstein I, Holinski-Feder E, Sala P, Gareth Evans D, et al. Cancer risk and survival in carriers by gene and gender up to 75 years of age: a report from the Prospective Lynch Syndrome Database. Gut 2018;67:1306–16.

[3] Valo S, Kaur S, Ristimäki A, Renkonen-Sinisalo L, Järvinen H, Mecklin J-P, et al. DNA hypermethylation appears early and shows increased frequency with dysplasia in Lynch syndrome-associated colorectal adenomas and carcinomas. Clin Epigenetics 2015;7:1–13.

[4] Campbell BB, Light N, Fabrizio D, Zatzman M, Fuligni F, de Borja R, et al. Comprehensive Analysis of Hypermutation in Human Cancer. Cell 2017;171:1042–56.e10.

[5] Allory Y, Beukers W, Sagrera A, Flández M, Marqués M, Márquez M, et al. Telomerase reverse transcriptase promoter mutations in bladder cancer: high frequency across stages, detection in urine, and lack of association with outcome. Eur Urol 2014;65:360–6.

[6] Bell RJA, Rube HT, Kreig A, Mancini A, Fouse SD, Nagarajan RP, et al. Cancer. The transcription factor GABP selectively binds and activates the mutant TERT promoter in cancer. Science 2015;348:1036–9.

[7] Borah S, Xi L, Zaug AJ, Powell NM, Dancik GM, Cohen SB, et al. Cancer. TERT promoter mutations and telomerase reactivation in urothelial cancer. Science 2015;347:1006–10.

[8] Comprehensive Genomic Profiling of Upper-tract and Bladder Urothelial Carcinoma. European Urology Focus 2021;7:1339–46.

[9] Donahue TF, Bagrodia A, Audenet F, Donoghue MTA, Cha EK, Sfakianos JP, et al. Genomic Characterization of Upper-Tract Urothelial Carcinoma in Patients With Lynch Syndrome. JCO Precis Oncol 2018;2018. https://doi.org/10.1200/PO.17.00143.

[10] Joost P, Therkildsen C, Dominguez-Valentin M, Jönsson M, Nilbert M. Urinary tract cancer in Lynch syndrome; Increased risk in carriers of MSH2 mutations. Urology 2015;86:1212–7.

[11] Nikkola J, Ryyppö L, Vuorinen J, Kallio H, Selin H, Jämsä P, et al. Sensitive Detection of Urothelial Cancer via High-volume Urine DNA Analysis. Eur Urol 2024. 10.1016/j.eururo.2024.10.014.

[12] Fujii Y, Sato Y, Suzuki H, Kakiuchi N, Yoshizato T, Lenis AT, et al. Molecular classification and diagnostics of upper urinary tract urothelial carcinoma. Cancer Cell 2021;39:793–809.e8.

[13] Heaphy CM, de Wilde RF, Jiao Y, Klein AP, Edil BH, Shi C, et al. Altered telomeres in tumors with ATRX and DAXX mutations. Science 2011;333:425.

[14] Suryo Rahmanto Y, Jung J-G, Wu R-C, Kobayashi Y, Heaphy CM, Meeker AK, et al. Inactivating ARID1A Tumor Suppressor Enhances TERT Transcription and Maintains Telomere Length in Cancer Cells. J Biol Chem 2016;291:9690–9.

[15] Cercek A, Lumish M, Sinopoli J, Weiss J, Shia J, Lamendola-Essel M, et al. PD-1 Blockade in Mismatch Repair-Deficient, Locally Advanced Rectal Cancer. N Engl J Med 2022;386:2363–76.

