## Supplement for "Constrained hypermutation and absence of TERT promoter mutations in Lynch syndrome-associated urothelial cancer"

### Supplementary Methods

**LS-UC tissue analysis**

A list of patients with a Lynch syndrome diagnosis was obtained from the Finnish Lynch Syndrome Registry, and all formalin-fixed, paraffin-embedded (FFPE) urothelial cancer tissue samples available in five Finnish biobanks were collected. If a patient had tissue available from multiple surgeries or from multiple lesions during one surgery, all such samples were analyzed (but only one sample per tumor lesion). All available slides per case were reviewed by a pathologist and re-classified according to WHO Classification of Tumours 5th edition. Representative cancerous areas and adjacent normal tissue (if available) on the 10 μm slides were identified and macrodissected using sterile scalpels, scraping tissue into 1.5ml Eppendorf tubes. DNA was extracted using the QIAamp DNA FFPE Advanced UNG Kit from the macrodissected FFPE tissue, eluted in 17-40 μl of ATE buffer and stored in -20C freezer. Immunohistochemical staining for MMR proteins used 4 μm slides. The slides were stained with antibodies for MLH1 (clone ES05, dilution 1:50, Novocastra, Leica Biosystems, Nussloch, Germany), MSH2 (FE-11, 1:50, Calbiochem, La Jolla, CA) and MSH6 (EPR3945, 1:150; Epitomics, Burlingame, CA) using a BOND-III Stainer and BOND Epitope Retrieval Solution 2 detection kit (Leica Biosystems). Based on nuclear staining in neoplastic cells, MMR protein expression was classified as lost or retained.

**Sporadic UC urine cohort**

Patients were recruited to a prospective urine collection study between January 2021 and January 2024 at the Tampere University Hospital. Patients with a positive diagnostic cystoscopy, prior diagnosis of UC, or a cystoscopy referral for primary UC evaluation (due to hematuria or imaging findings) were invited to enroll. The study was conducted in accordance with the Declaration of Helsinki, and the protocol was approved by the Regional Ethics Committee of Tampere University Hospital (identification code R21088B). All patients provided written informed consent before study participation.

Patients were instructed to collect 100 mL of first void urine (with a recommended minimum of 2 hours since last urination) into a plastic cup at the Tampere University Hospital Urology Clinic or a sample bag at home. Patients added 5 mL of Streck Urine DNA preservative to the sample immediately after collection to ensure sample integrity at room temperature. Home urine samples were transported to Tampere University via standard mail.

**Library construction and DNA sequencing**

Illumina DNA sequencing libraries were constructed from 24 - 100 ng of DNA using the Twist Enzymatic Fragmentation Library Prep Kit 2.0 with 7.5 min fragmentation time and following the manufacturer's recommended protocol. Sequencing libraries were multiplexed into pools, and target capture was carried out using custom IDT xGen hybridization capture panels and xGen™ Hybridization and Wash v2 Kit (IDT). The library pools were sequenced on an Illumina NovaSeq 6000 instrument using 2x150 bp paired end sequencing with v1.5 flow cells.

**Somatic mutation analysis**

To identify somatic mutations (base substitutions and indels) in FFPE tumor tissue samples, we searched for variants with ≥ 7 supporting reads, allele fraction ≥ 5%, and allele fraction ≥ 20 times the background error rate in negative control samples. For mutations in urine tumor DNA samples, an allele fraction ≥ 1% was required instead, given the lower rates of DNA damage in fresh nuclear DNA extracted from urine cell pellets. For FFPE tissue samples sequenced using the UroScout assay, we quantified the position and allele specific background error rate based on negative control urine samples from 24 healthy volunteers (median age 39 years). For FFPE tissue samples sequenced with the whole exome panel, we quantified background error rates based on a set of 20 adjacent normal FFPE tissue samples. Mutation calling thresholds were halved for a curated set of 38 highly recurrent urothelial cancer mutations (**Supplementary Table 4**). We required that the average distance of the mutant allele from the nearest read end was ≥ 14 bases, and the average mapping quality of mutant allele carrying reads was ≥ 10. Variants found in the gnomAD v3.0 germline variant database with a population frequency ≥ 0.5% were discarded [16]. Protein-level consequences of mutations were predicted using ANNOVAR [17].

**Quantification of sample cancer fraction**

We defined the cancer fraction of a tissue or urine sample as the fraction of genome equivalents in the sample originating from cancer cells, and used the formula $C = 2 / (1 / F + 1)$ to solve cancer fraction $C$ based on the allele fraction $F$ of each somatic mutation detected in the sample, and selected the highest cancer fraction estimate. This formula conservatively assumes loss-of-heterozygosity through deletion of the non-mutant allele.

**References**


[16] Karczewski KJ, Francioli LC, Tiao G, Cummings BB, Alföldi J, Wang Q, et al. The mutational constraint spectrum quantified from variation in 141,456 humans. Nature 2020;581:434–43.

[17] Wang K, Li M, Hakonarson H. ANNOVAR: functional annotation of genetic variants from high-throughput sequencing data. Nucleic Acids Res 2010;38:e164.

### Supplementary Figures

Patient 29
Bladder cancer
MSH6

Patient 19
UTUC
MSH2

Patient 23
UTUC
MLH1

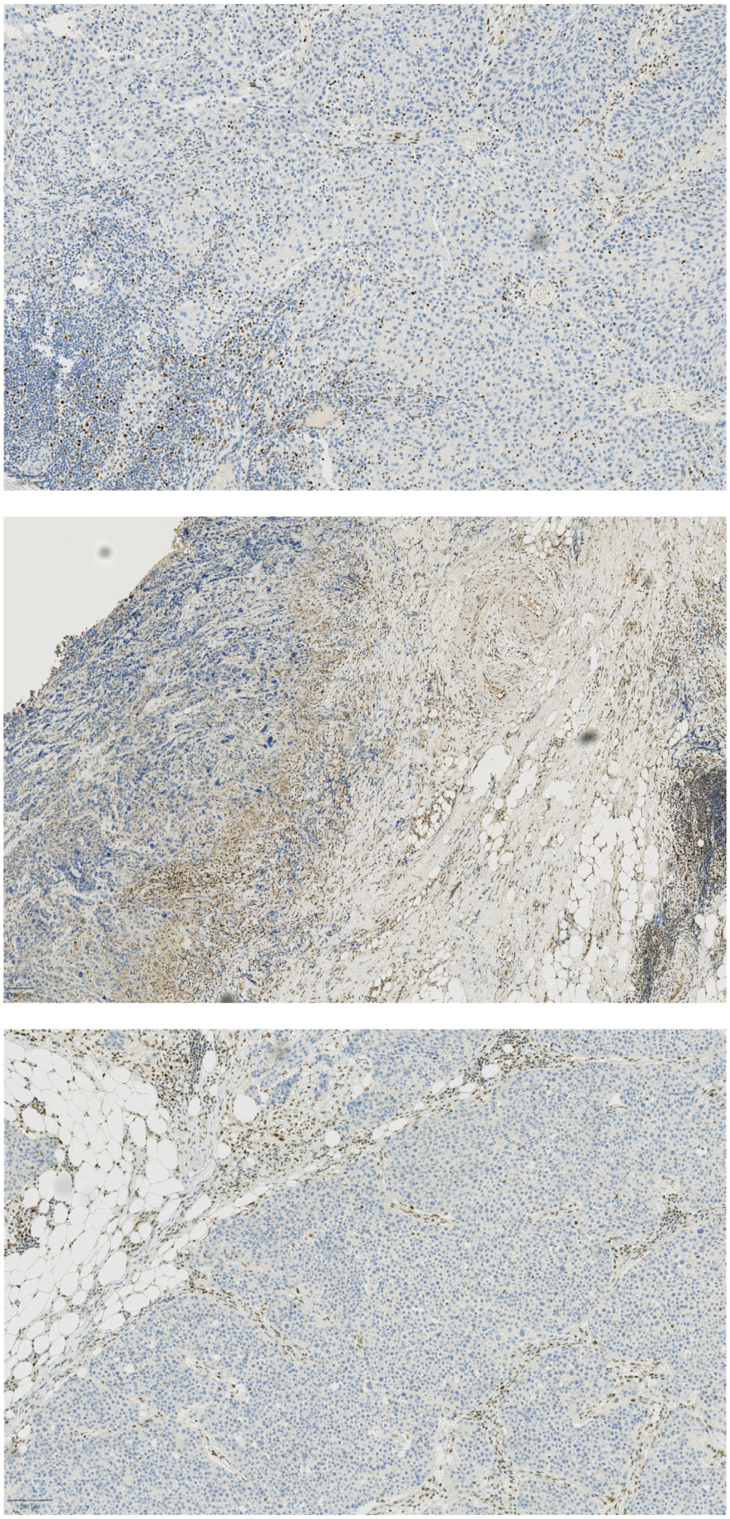

**Supplementary Figure 1.** Immunohistochemical confirmation of DNA mismatch repair gene inactivation in Lynch syndrome-associated urothelial cancer tissue in our cohort. Three representative cases are shown (one each for MSH2, MSH6, and MLH1).

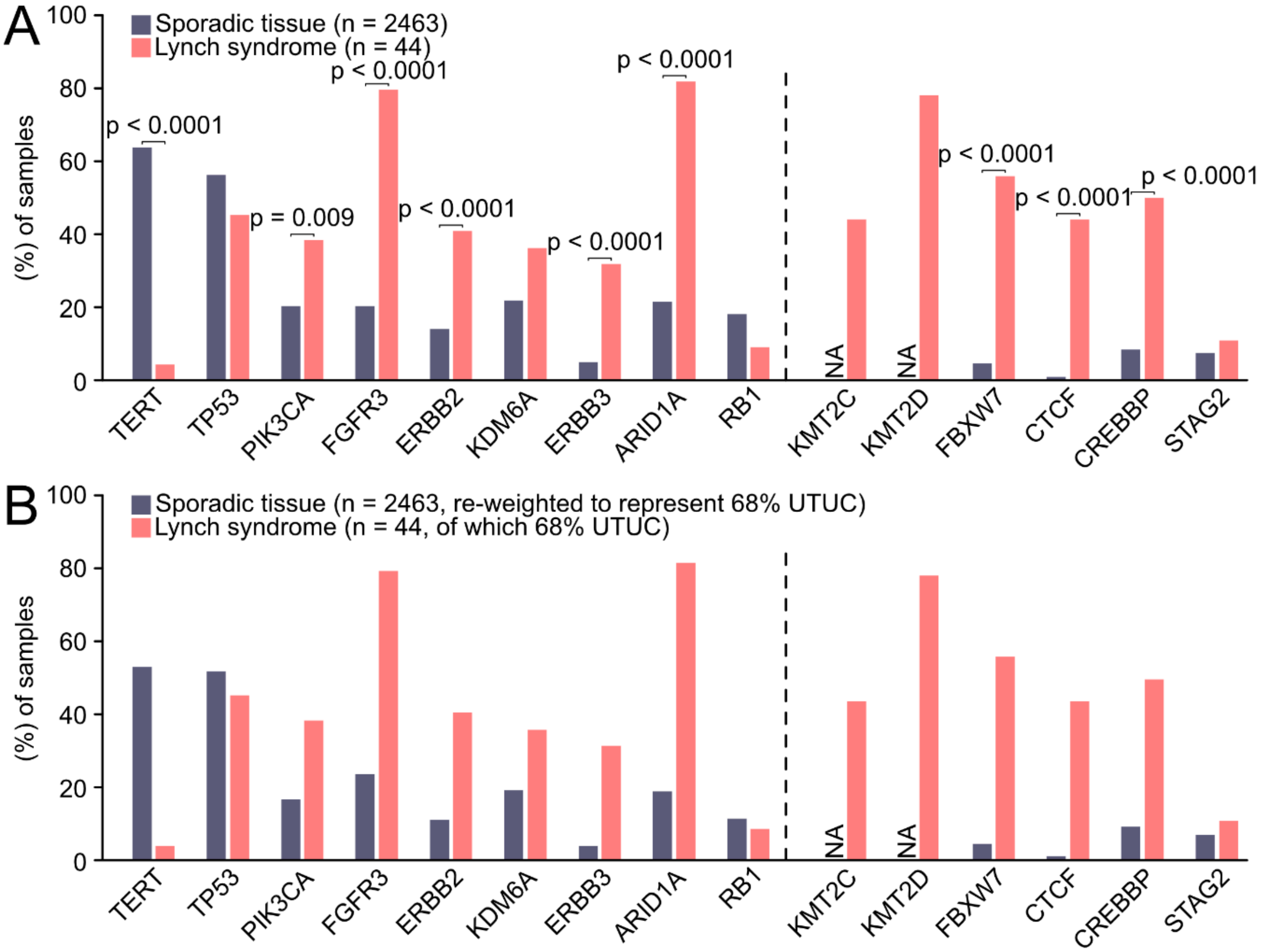


**Supplementary Figure 2.** Comparison of driver gene mutation frequencies in our Lynch syndrome-associated urothelial cancer tissue cohort analysed with the UroScout assay (n = 44 samples, left-side of the dashed line), with whole exome sequencing (n = 18 samples, right-side of the dashed line) and a large sporadic urothelial cancer (UC) tissue cohort by Necchi et al [8]. (**a**) Direct cohort-to-cohort comparison. (**b**) Adjusted comparison where the driver gene mutation frequency in the sporadic UC cohort is weighted to normalize for the different rate of upper tract urothelial cancer (UTUC) and bladder cancer in the LS-UC and sporadic UC cohorts.

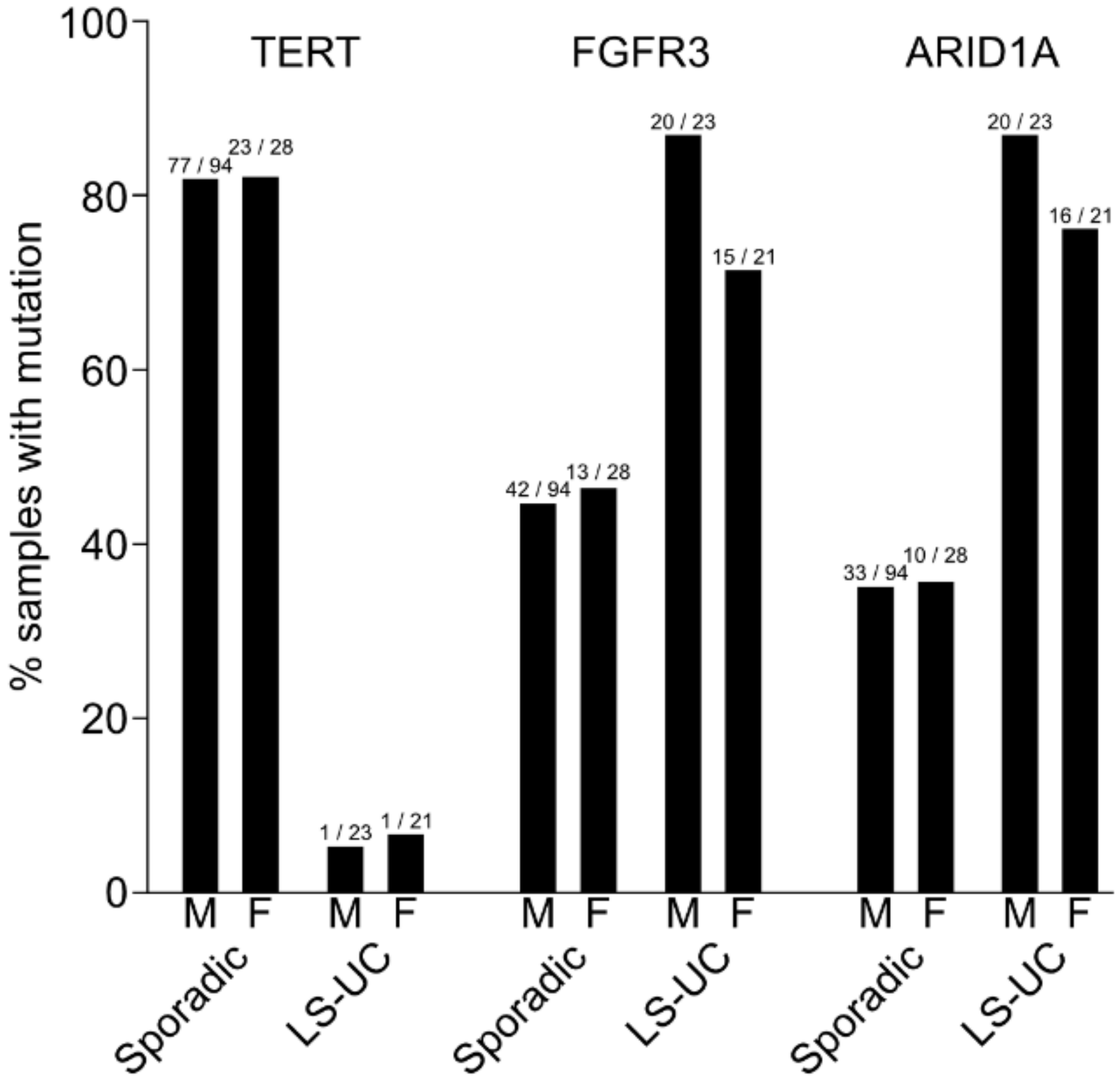


**Supplementary Figure 3.** Mutation frequency of *TERT*, *FGFR3*, and *ARID1A* in sporadic urothelial cancers (UC) (n=122) and Lynch syndrome-associated urothelial cancers (LS-UC) in patients of different sex (M = male, F = female). While sporadic UC is more common in males, LS-UC incidence is more equal between the sexes. However, no statistically significant association between patient sex and mutation frequency was identified for any of the three genes, indicating the patient sex does not explain the differences in gene mutation frequency between sporadic and LS-UC.

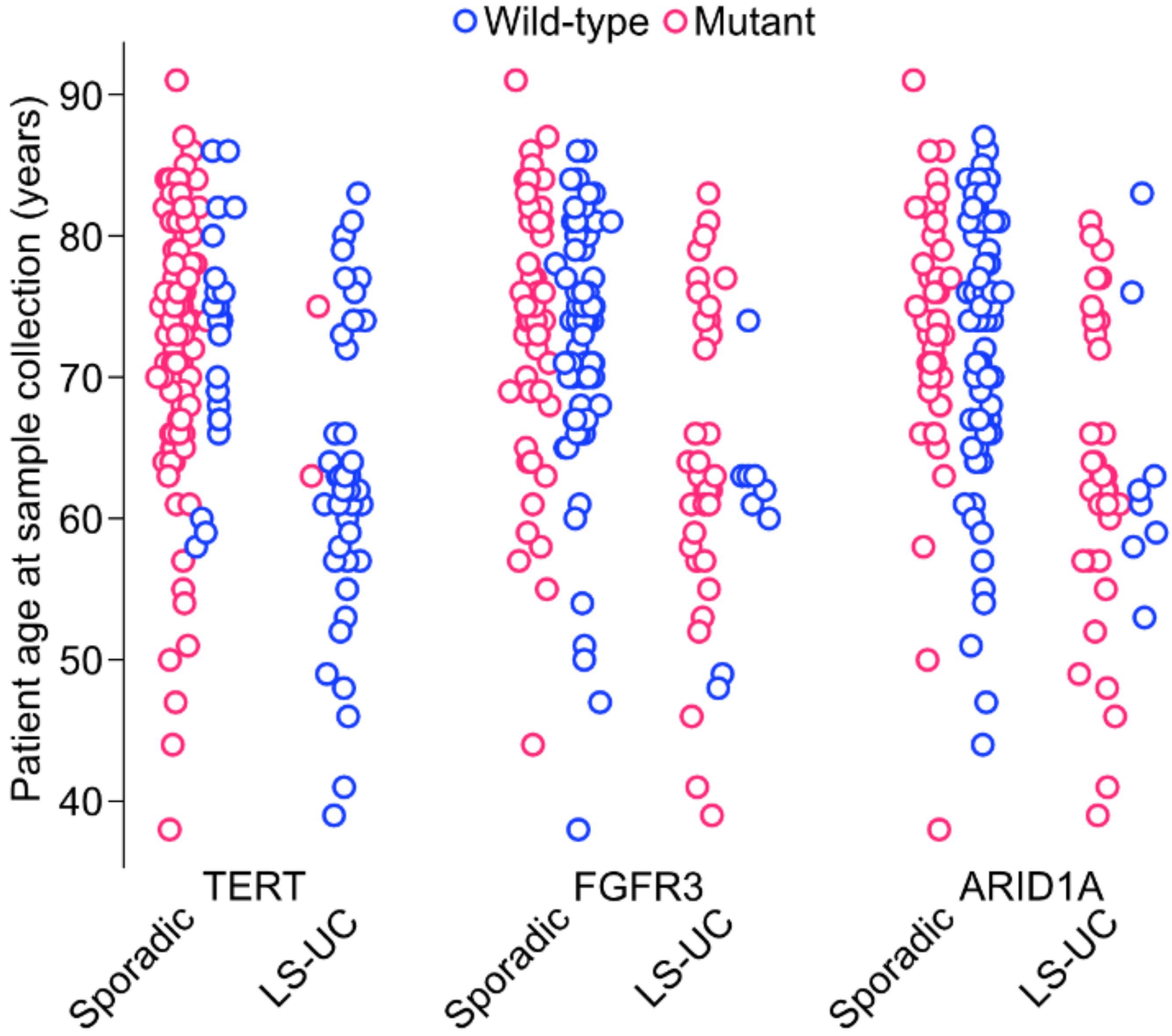


**Supplementary Figure 4.** Mutation frequency of *TERT*, *FGFR3*, and *ARID1A* in sporadic urothelial cancers (UC) (n=122) and Lynch syndrome-associated urothelial cancers (LS-UC) in patients of different ages. Each dot is a tumor, colored by mutation status. While LS-UC tumors are typically diagnosed at a younger age than sporadic UC, this visualization shows that patient age does not explain the difference in driver gene mutation frequencies between sporadic and LS-UC.

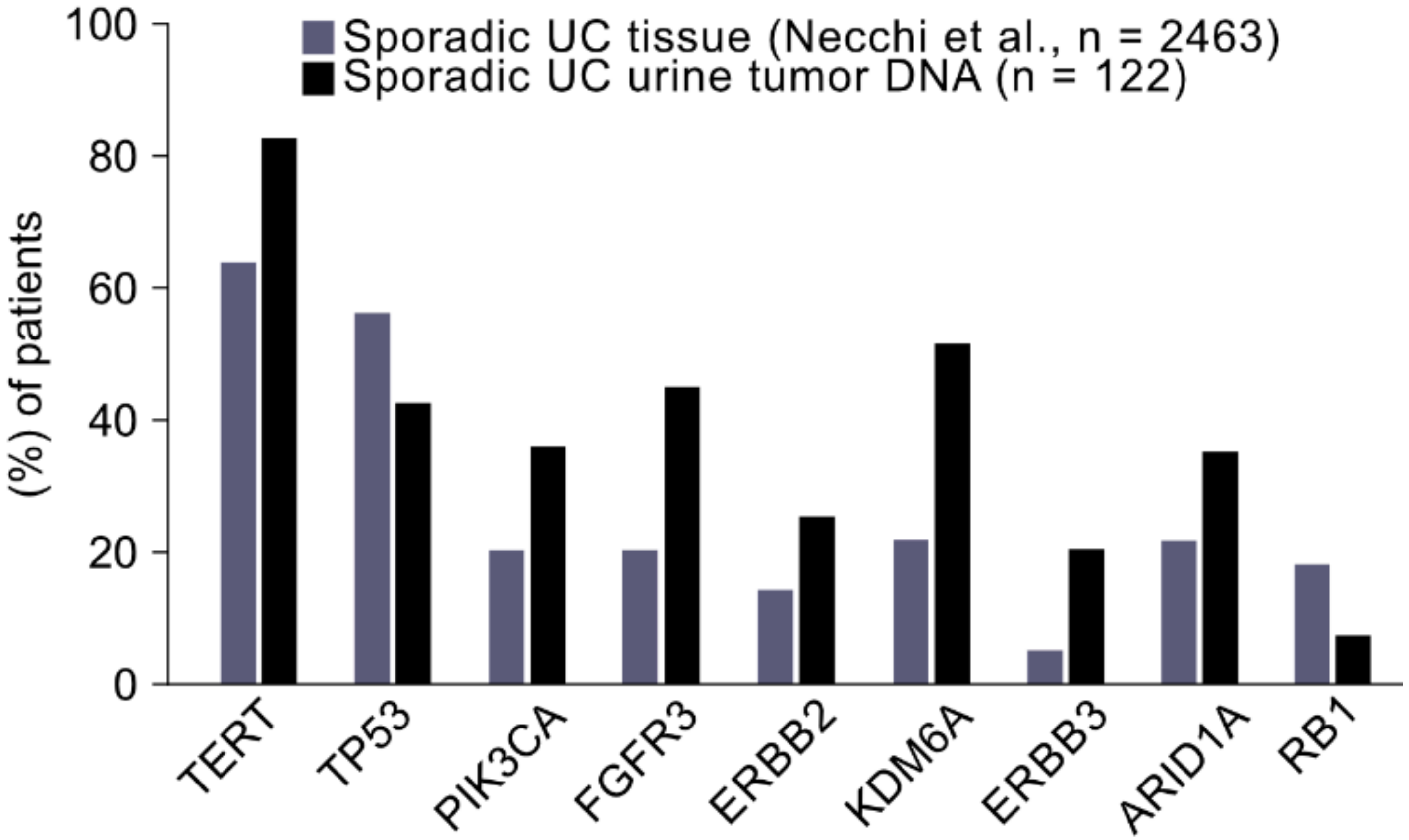


**Supplementary Figure 5.** Comparison of driver gene mutation frequency in our sporadic all-comers urothelial cancer (UC) urine sample cohort (n = 122 samples) and the Necchi et al. sporadic UC tissue cohort [8]. The Necchi et al. sporadic UC tissue cohort included only high grade and metastatic UCs, explaining the higher rate of *TP53* and *RB1* mutations that are correlated with aggressive disease, and the reduced rate of *FGFR3* mutations that are correlated with less invasive papillary tumors.

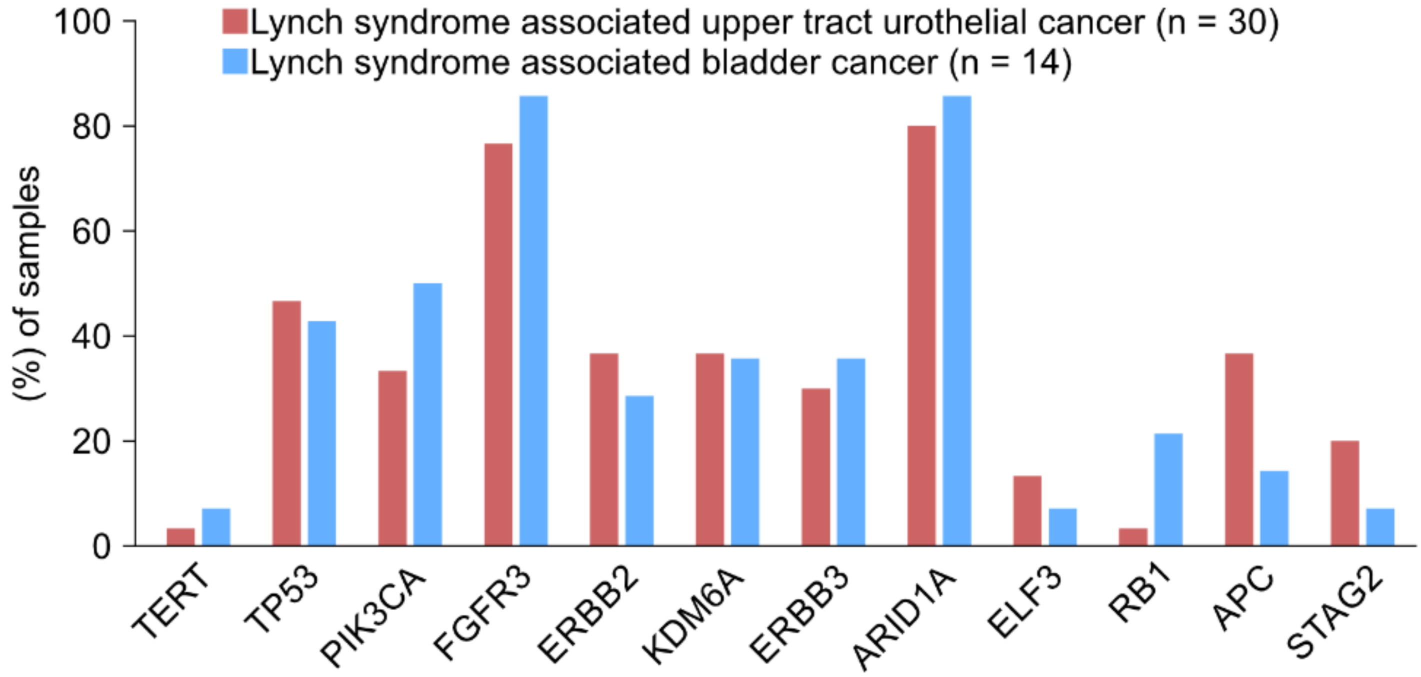


**Supplementary Figure 6.** Comparison of driver gene mutation frequency between Lynch syndrome-associated bladder cancers (n = 14) and upper tract urothelial cancers (n = 30) in our cohort.

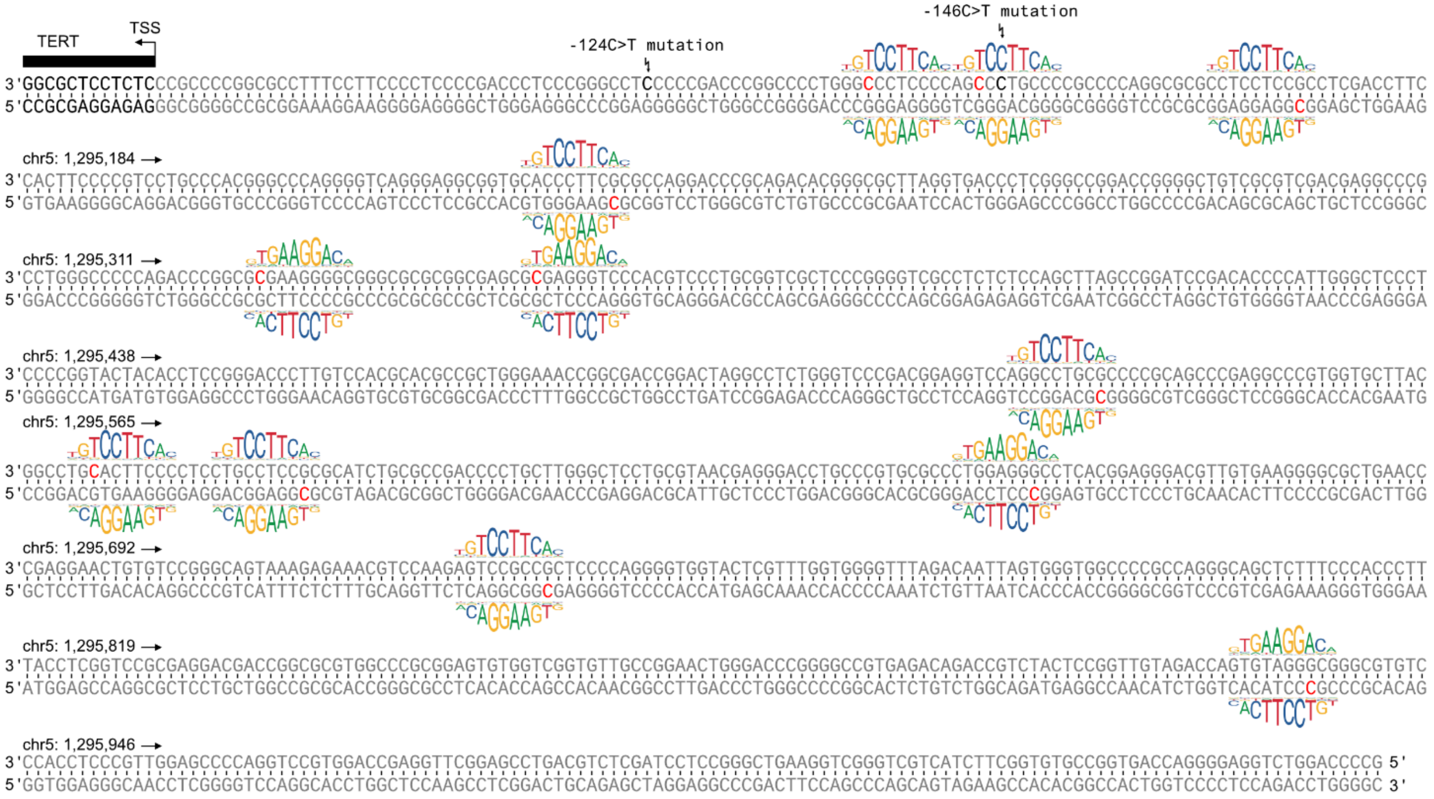


**Supplementary Figure 7.** Visualization of potential sites within the *TERT* promoter region where a CG>TG mutation can produce a DNA sequence approximating the GABP binding motif. The double-stranded DNA sequence up to 1000 bp upstream of the *TERT* transcription start site is shown. GABP binding motifs (JASPAR identifier MA0062.3) are shown at positions where a CG>TG mutation can produce a DNA sequence with a $p < 0.001$ motif match score (calculated using the PWMScan software). The two most common TERT promoter mutations are also indicated (-124C>T and -146C>T, neither one is a CG>TG type mutation). CG>TG mutations cannot produce the most important 4-base 5’-GGAA-3’ sequence in the center of the GABP binding motif due to the lack of a TG dinucleotide in the core motif. TG dinucleotides are only found in the less conserved flanks of the GABP binding motif (see Figure 1f). No actual CG>TG somatic mutations reinforcing these near-match GABP binding motif sites were detected in our Lynch syndrome-associated urothelial cancer cohort.
